# Role of Bassoon-mediated active zone integrity at different types of brain synapses for brain activity and cortex-dependent memory formation

**DOI:** 10.1101/2025.11.24.690204

**Authors:** Anja M. Oelschlegel, Horst Schicknick, Anil Annamneedi, Carolina Montenegro-Venegas, Anna Fejtová, Eckart D. Gundelfinger, Jürgen Goldschmidt, Wolfgang Tischmeyer

## Abstract

**Background:** The properly controlled release of neurotransmitter at presynaptic active zones is crucial for brain function and performance. Bassoon is a major scaffolding protein involved in the organization of neurotransmitter release sites at excitatory, inhibitory and modulatory brain synapses. Global deficiency of functional Bassoon causes severe neurological conditions including disturbed patterns of brain activity and early-onset epilepsy. To distinguish the contribution of different types of synapses to this phenotype we generated conditional knockout (cKO) mice lacking the *Bsn* gene in (i) GABAergic interneurons expressing Cre recombinase under the control of the *Dlx5/6* regulatory elements (*Bsn^Dlx5/6^* cKO), (ii) glutamatergic forebrain neurons expressing Cre under the *Emx1* promoter (*Bsn^Emx1^* cKO), and (iii) dopaminergic neurons expressing *DAT-*driven Cre (*Bsn^DAT^*).

**Methods:** Single-photon emission computed tomography (SPECT) imaging of cerebral blood flow (CBF) was employed to assess *in vivo* brain-wide activation patterns in the cKO mice and corresponding control animals with wildtype *Bsn* genes. A cortex-dependent learning task to discriminate frequency-modulated tones was then used to evaluate the cognitive abilities of the different cKO lines.

**Results:** Marked reduction of brain activity was found in various cortical areas and the basolateral amygdala of *Bsn^Dlx5/6^* cKO mice, while patches of increased activity were detected in the dorsal striatum. *Bsn^Emx1^* cKO mice, in contrast, display increased brain activity in many cortical areas. Only minor changes in CBF were detected in *Bsn^DAT^* cKO mice. Concerning auditory discrimination learning *Bsn^Dlx5/6^* cKO mice were severely impaired, although they responded to stimuli normally. On the other hand, *Bsn^Emx1^* cKO mice acquired the task more efficiently reaching maximum performance levels faster than control animals. Surprisingly, *Bsn^DAT^* cKO mice did not differ in their behavior from the control group.

**Conclusions:** Our data suggest that absence of Bassoon from presynapses of GABAergic interneurons expressing *Dlx5/6* gene during development results in severe neurological symptoms and associated dysfunctions. Instead, network changes associated with Bassoon deficiency at glutamate release sites of excitatory forebrain neurons, even seem to have an enhancing effect on learning.

**Graphical Abstract:** 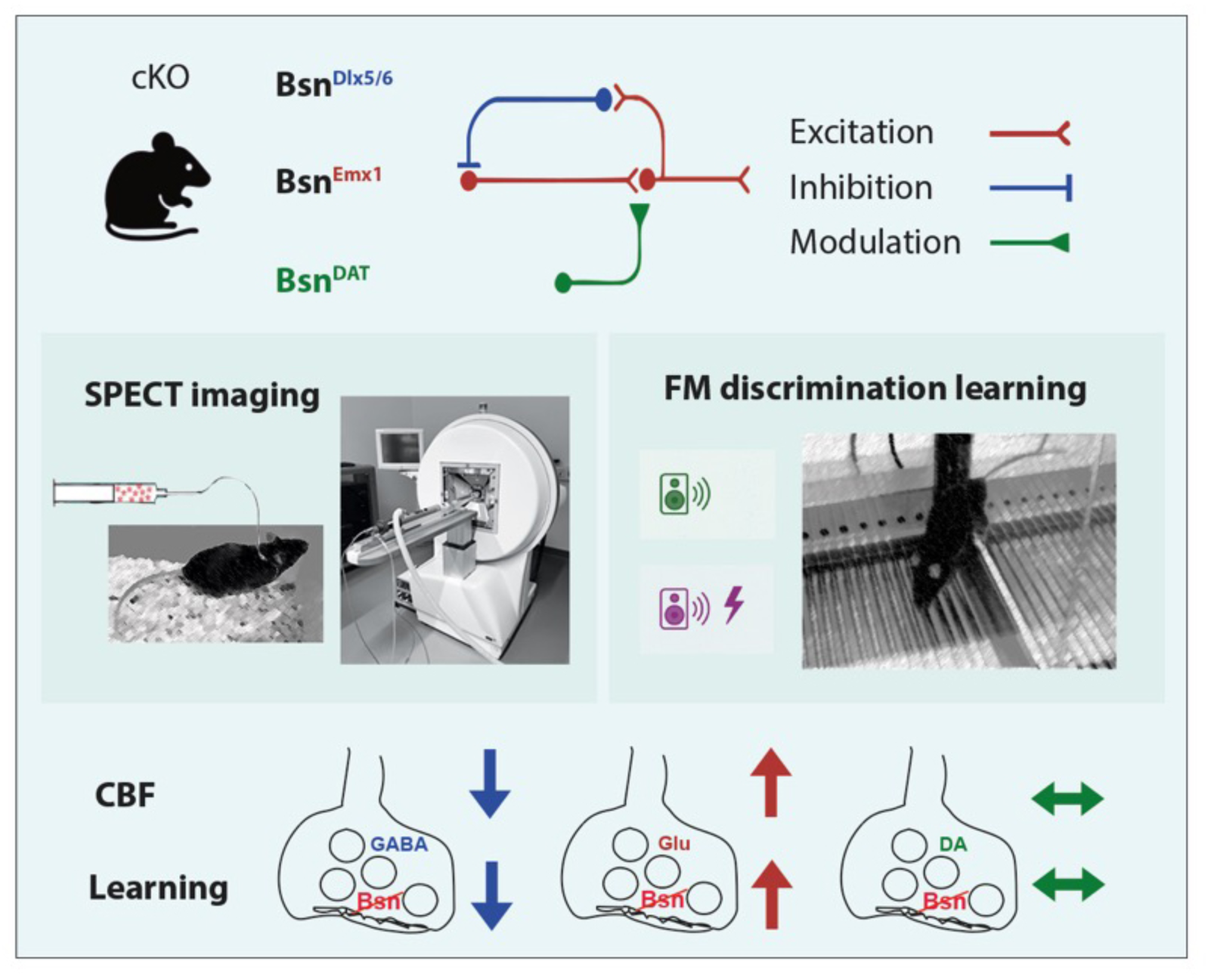

## Background

Regulation of neurotransmitter release is a major determinant of brain plasticity and accordingly plays an important role in processes underlying memory formation (*e.g*. [1]). The presynaptic scaffolding protein Bassoon, which is present at active zones of neurotransmitter release of both excitatory and inhibitory brain synapses [2, 3], is an essential organizer of the release apparatus. The presence of Bassoon has also been reported for release sites from dopaminergic neurons [4–8]. Bassoon has multiple functions in the developmental assembly, molecular organization and functional plasticity of the cytomatrix at the active zone of neurotransmitter release [9]. Thus, it is involved in the localization of P/Q-type voltage-gated calcium channels in the active zone membrane [10], in the control of presynaptic proteostasis [11–13] and in the refilling of synaptic vesicle release sites [14, 15]. In addition, Bassoon, together with its paralogue Piccolo, regulates the distribution of the chromatin-modulating protein CtBP1 between neuronal nuclei and synapses [16, 17], suggesting that it may in this way affect gene expression during neuronal maturation and plasticity.

Global lack or dysfunction of Bassoon in the CNS has severe consequences. In mice, these include epilepsy with rapidly generalizing seizures [18–21]; abnormal brain enlargement combined with cytoarchitectural abnormalities and disturbed metabolism of glutamatergic metabolites in the cortex [22, 23] and increased levels of BDNF [24, 25]; as well as sensory deficits because of impaired anchoring of presynaptic ribbons at retinal photoreceptor and inner hair cell synapses [26–29]. In human patients, mutations or dysregulation of the *BSN* gene have been associated with a variety of neurodevelopmental and neurodegenerative brain disorders. These include Rolandic forms of epilepsy in children [30–32] that can be associated with intellectual disability [33], as well as schizophrenia [34], Tourette syndrome [35] or early-onset parkinsonism [36]. Moreover, mutations in the *BSN* gene have been associated with progressive supranuclear palsy-like syndrome [37, 38] by serving as potential seed for tau aggregation [39]. The presence of both active zone scaffolders, Bassoon and Piccolo, in the synaptic proteomes is regulated by processes of synaptic plasticity. Significant down-regulation of both proteins occurs during homeostatic adaptation of synaptic strength in primary cultured hippocampal neurons [40] as well as in the dorsal hippocampus after contextual fear learning [41] or in the striatum and frontal cortex upon auditory discrimination learning [42]. In contrast, neonatal irradiation was reported to induce significant up-regulation of the two proteins in cortex and hippocampus [43].

In an approach to disclose Bassoon functions in different neurons and at different types of synapses we have generated conditional knockout (cKO) mice by inactivating the *Bsn* gene in a subset of inhibitory – mainly parvalbumin expressing (PV+) – neurons (*Bsn^Dlx5/6^* cKO), in forebrain excitatory neurons (*Bsn^Emx1^* cKO), and in dopaminergic neurons (*Bsn^DAT^* cKO) [19, 44, 45]. *Bsn^Dlx5/6^* cKO mice are epileptic [19] and have major learning deficits in various paradigms [45, 46]. *Bsn^Emx1^* cKO mice show only relatively mild epilepsy [19], impaired maturation of excitatory synapses and neurons combined with maintenance of juvenile-like plasticity in the dentate gyrus and with enhanced performance, *e.g.*, in contextual fear learning, spatial pattern separation and novel object location detection tasks [44]. *Bsn^DAT^* cKO mice show only very mild phenotypes, altogether; they display slight changes in anxiety and safety learning [45].

Here, we asked, how conditional removal of Bassoon from different types of synapses affects signal processing at network or system levels and studied brain-wide activation patterns and cortex-dependent learning behavior in different *Bsn* cKO mutants.

For brain activity assessment, we employed single-photon emission computed tomography (SPECT) to image *in vivo* the spatial patterns of cerebral blood flow (CBF), a well-established proxy for neural activity, in *Bsn* cKO mice and corresponding controls. CBF-SPECT allows to map brain-wide activation patterns in awake, freely moving animals [47]. This approach is similar in rationale to ^18^F-2-fluoro-2-deoxyglucose positron emission computed tomography widely used in humans and animals, but it provides higher spatial resolution in rodents [47].

As the different *Bsn* cKO mice showed clear differences in the CBF within cortical areas, we tested another set of animals in a complex, cortex-dependent learning task to correlate disturbances of brain activity patterns with corresponding behavioral performances. Frequency-modulated tone (FM) discrimination learning is a demanding Go/No-Go task to discriminate short tone sequences of rising and falling frequencies to avoid electrical foot shocks in a shuttle box [48, 49]. Given that cortical dopamine and excitatory/inhibitory balance may determine the efficiency of FM discrimination learning [50–52], we used this paradigm to assess the learning capabilities of the different *Bsn* cKOs. While *Bsn^DAT^* cKO mice displayed no significant behavioral changes and only minor CBF changes compared to control animals, mice lacking Bassoon in excitatory versus inhibitory neurons of the forebrain exhibited significant and quite contrasting changes in learning behavior and brain activity patterns.

## Materials and Methods

### Animals

Generation of mice with *loxP* sites flanking exon 2 of the *Bsn* gene encoding the presynaptic scaffolding protein Bassoon (*Bsn^lx/lx^ / Bsn^tm1.1Arte^*) as well as of the conditional knockout lines *Bsn^Dlx5/6^*, deficient for Bassoon in a subset of inhibitory interneurons of the forebrain upon expression of Cre recombinase under the control of Dlx5/6 regulatory elements [53, 54], and *Bsn^Emx1^*, lacking expression of Bassoon in excitatory forebrain neurons upon expressing Cre recombinase under the control of the *Emx1* regulatory sequences [55], has been described previously [19, 44]. Mice with conditional deletion of the *Bsn* gene in dopaminergic neurons (*Bsn^DAT^*) were generated by crossing *Bsn^lx/lx^* mice with animals expressing Cre recombinase under the transcriptional control of the endogenous DAT promoter (*Slc6a3^tm1.1(cre)BKmn^*) [45, 56]. Three genetic mouse variants with wild-type genotype regarding the *Bsn* gene served as controls: *Bsn^lx/lx^* mice (w/o Cre expression) as controls for *Bsn^Emx1^* and *Bsn^Dlx5/6^*; *Bsn*^+/+^ x *Slc6a3^tm1.1(cre)BKmn^* mice as reference for *Bsn^DAT^*, and *Bsn^+/+^* x *Emx1^tm1(cre)Krj^* as a further reference for *Bsn^Emx1^* (Additional file 1). Mice were housed under standard conditions on a 12h/12h light-dark cycle. Food and water were provided *ad libitum*. All *Bassoon* mutant lines were bred on a C57BL/6N genetic background at the LIN animal facility.

**Animal licenses:** All animal experiments were conducted in accordance with the European and German regulations for animal experiments and were approved by the Landesverwaltungsamt Sachsen-Anhalt (Numbers: 42502-2-988 LIN, 42502-2-1303 LIN, and 42502-2-1589 LIN).

### SPECT-imaging of CBF

Between 8-15 weeks-old mice of both sexes (*Bsn^Dlx5/6^* cKO n=10, ctrl. n=9; *Bsn^Emx1^* cKO n= 9, ctrl. n=8; *Bsn^DAT^*cKO n=12, ctrl. n=12) were implanted with an intravenous catheter into the right external jugular vein [47, 57]. They were given at least one day for recovery before imaging. The catheter was subcutaneously tunneled and exited in the neck region between the scapulae. Before tracer-injection the catheter was connected via a 0.9% saline-filled plastic tube 60 cm in length to a ^99m^Technetium hexamethyl propylene amine oxime (^99m^Tc-HMPAO) filled syringe in a perfusion pump. Mice were able to freely move in a cage open at the top while being i.v. injected through the plastic tube and catheter with freshly prepared ^99m^Tc-HMPAO [47]. Doses of on average 135 MBq (+/- 14) were injected in 330 µl within 15 min. After injection, mice were anesthetized with isoflurane (3%, 900 ml/min O_2_) and scanned under isoflurane anesthesia (1.2-1.5%, 900 ml/min O_2_) in a NanoSPECT/CT^TM^ scanner (Mediso, Hungary) with an acquisition time of 60 min.

SPECT scans were made using ten-pinhole mouse brain apertures with 1.0 mm pinhole diameters providing a nominal spatial resolution < 0.7 mm. Photopeaks were set to the default values of the NanoSPECT/CT for ^99m^Tc (140 keV ± 5%). SPECT images were reconstructed at an isotropic voxel size of 250 µm using the manufactureŕs software (HiSPECT^TM^, SCIVIS, Germany). SPECT scans were accompanied by two co-registered CT scans, one before and one after the SPECT scan, to control for motion artefacts (none detected). CT scans were made with an acquisition time of 90 sec at 45 kVp, 177 μA, with 180 projections, 500 ms per projection, and 96 μm isotropic spatial resolution, reconstructed with the manufactureŕs software (InVivoScope 1.43) at isotropic voxel-sizes of 100 µm.

SPECT scans were aligned to an anatomical reference MR using the co-registered CT scans in an automated alignment workflow [58]. After alignment, SPECT data were converted to TIFF-files, brain data were cut out from the head scans using a brain mask and global-mean normalized in ImageJ. An unpaired two-tailed heteroscedastic voxel-wise t-test was made in MATLAB (version R2017b (9.3.0.713579) 64-bit, Mathworks, USA). T-test result-files (uncorrected for multiple comparisons) as well as group-mean and difference-files were converted to DICOM and overlaid in the OsiriX^TM^ software (version 5.9.1, Pixmeo, Switzerland) on an anatomical reference MR from the literature [59]. Selected sections were exported and arranged as a composite figure in Photoshop (version 24, Adobe, USA).

### FM discrimination paradigm

None of the three *Bsn* cKO mouse lines had shown alterations in startle magnitude when compared to their control littermates [45] indicating that they are not impaired in their hearing capabilities. Behavioral training and statistics were performed as described previously [60, 61]. Essentially, 10-19 weeks-old male mice were trained once per day, for 16 sessions in total, on a foot-shock reinforced shuttle box avoidance Go/No-Go procedure to discriminate the directions of frequency modulations. During a 3-min adaptation period preceding each training session mice were allowed to habituate to the training chamber. During sessions, animals were trained to discriminate between conditioned stimuli (CSs) consisting of sequences (250-ms tone, 250-ms pause) of an ascending (4–8 kHz, CS+) and a descending (8–4 kHz, CS-) FM. A training session of 25 minutes consisted of 30 presentations of each, CS+ and CS-, in a pseudo-randomized order. The mean inter-trial interval was 15 s. To avoid electrical foot shock, mice had to cross the hurdle of the shuttle box within 6 s of CS+ and to suppress this response within 6 s of CS- presentations. Hurdle crossings within 6 s upon the onset of CS+ and CS- were regarded as correct conditioned responses (CR+) and false alarms (CR-), respectively. For each session, the numbers of CR+ and CR- were monitored and the relative frequencies of CR+ and CR- were calculated as percentage of trials with presentations of CS+ and CS-, respectively. To quantify the discrimination performance, the discrimination rate (*D*), *i.e.*, the difference between the relative frequencies of CR+ and CR-, was calculated. To assess effects on learning the hurdle reaction in response to the FMs *per se*, the sum of CR+ and CR- (ΣCR) expressed as percent of the total number of trials, was calculated. To assess effects on sensory, motivational and motor systems, the avoidance latency (tCR+), *i.e.*, the time required to change the compartment during CR+, was recorded (with failed trials assigned a latency of 6 s, the maximum length of the CS). To assess general arousal and activity of the experimental animals, the pre-session activity (PSC), *i.e.*, the numbers of hurdle crossings during the 3-min adaptation period preceding each session, and the inter-trial activity (ITC), *i.e.*, the average number of hurdle crossings occurring per inter-trial interval of each session, were monitored. Behavioral data are presented as group means ± S.E.M. Data without index were expressed as group means per session. When linked to the index “block” (cf. inset of Figure 2A), a training session was subdivided into five blocks of trials, and data were expressed as group means per trial block. Each trial block consisted of 12 consecutive trials, that is, six presentations of each CS+ and CS-. For statistical evaluation, StatView 5.0.1 (SAS) was used. Repeated-measures analysis of variance (RM-ANOVA), with training session or, where indicated (cf. inset of Fig. 2A), trial block serving as the repeated measures, was performed as indicated. Student’s two-tailed *t*-tests for paired or unpaired comparisons were used where appropriate. *P* values of <0.05 were considered as statistically significant.

## Results

### SPECT imaging of cerebral blood flow in the various *Bsn* cKO mutants

SPECT imaging of CBF in rodents makes use of the intravenously applied lipophilic tracer ^99m^Tc-HMPAO, which upon crossing the blood-brain barrier is rapidly converted to a hydrophilic compound that is trapped in the brain and shows no further redistribution (for review see ref. [47]). The ^99m^Tc distribution in the brain can be read out in anesthetized animals in the SPECT scanner and provides an integrated measure of blood flow in the awake brain during the timespan of injection.

Significant changes in the spatial patterns of baseline CBF were found in all groups with conditional *Bsn* ablations as compared to according control groups. In *Bsn^Dlx5/6^* cKO mice (Fig. 1A-F) marked decreases in mean CBF with values up to 25% were found in many cortical areas passing thresholds of significance in the anterior cingulate (Fig. 1A), primary motor cortex (Fig. 1C), auditory cortex (Fig. 1D) and visual cortex (Fig. 1F). CBF decreased also in the basolateral / basomedial region of the amygdala (Fig. 1D). Increased mean CBF was found in several patches within the dorsal striatum passing threshold of significance in one of the patches (Fig. 1B). In addition, significantly increased CBF was found in the periaqueductal gray (Fig. 1F).

**Figure 1.**
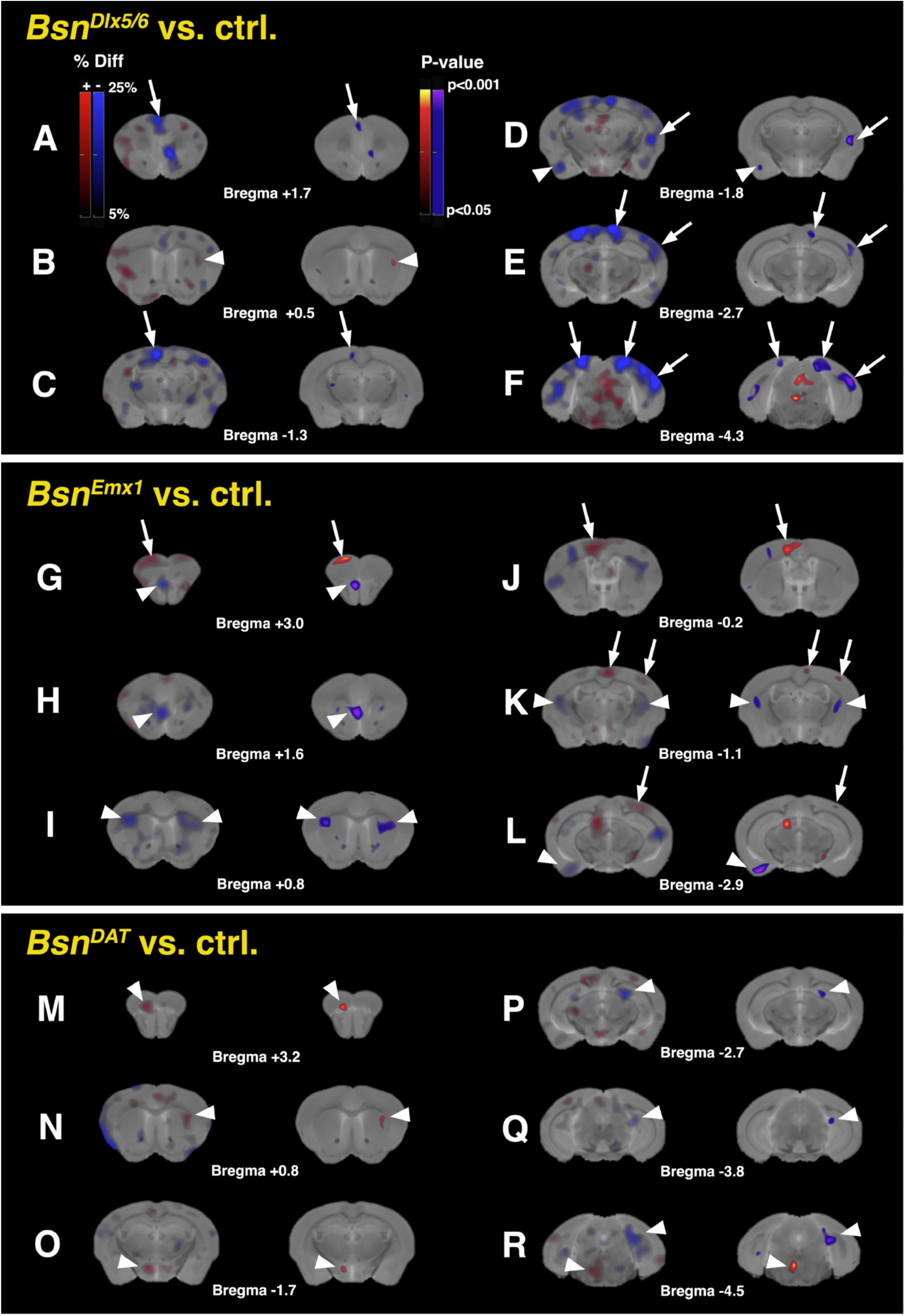
Spatial patterns of baseline CBF alterations in mice with conditional *Bsn* ablations versus corresponding controls. Shown are maps of percentage differences (% Diff) in group-mean intensity-normalized CBF and statistically significant differences (P-value) between cKO and corresponding control mice in rostro-caudal series of frontal sections at six different Bregma levels. Results from comparisons of conditional ablation in distinct GABAergic neurons (*Bsn^Dlx5/6^*) vs. controls are shown in the panels **A-F**, in forebrain glutamatergic neurons (*Bsn^Emx1^*) in panels **G-L**, and in dopaminergic neurons (*Bsn^DAT^*) in panels **M-R**. Within each group of panels, two images at the same Bregma level are shown with differences in mean CBF in the left and p-values in the right section. Warm colors indicate increased, cold colors decreased mean CBF in the mutants as compared to their corresponding controls. Note the strong decrease in CBF of up to 25% in cortical regions of the *Bsn^Dlx5/6^* cKO mice (arrows in **A, C, D, E, F**). Changes of opposite sign are found in cortical regions in the *Bsn^Emx1^* cKO mice (arrows in **G, J, K, L**). CBF decreases in the olfactory bulb (arrowhead in **G**) and in the dorsal peduncular cortex / dorsal tenia tecta region (arrowhead in **H**) of *Bsn^Emx1^* cKO mice and in parts of the amygdala in both *Bsn^Dlx5/6^* and *Bsn^Emx1^* mutants (arrowheads in **D, L**). In all three mutants CBF is altered in patches within the dorsal striatum (arrowheads in **B, I, K, N**). In the *Bsn^DAT^*cKO mice alterations in mean CBF are comparably small. Significant increases are found in orbitofrontal cortex (arrowhead in **M**), dorsomedial hypothalamus (arrowhead in **O**) and brainstem raphe (arrowhead in **R**), significant decreases in dentate gyrus (arrowheads in **P, Q**) and presubiculum (arrowhead in **R**). Animal numbers: *Bsn^Dlx5/6^* cKO n=10, ctrl. n=9; *Bsn^Emx1^*cKO n= 9, ctrl. n=8; *Bsn^DAT^* cKO n=12, ctrl. n=12.

**Figure 2.**
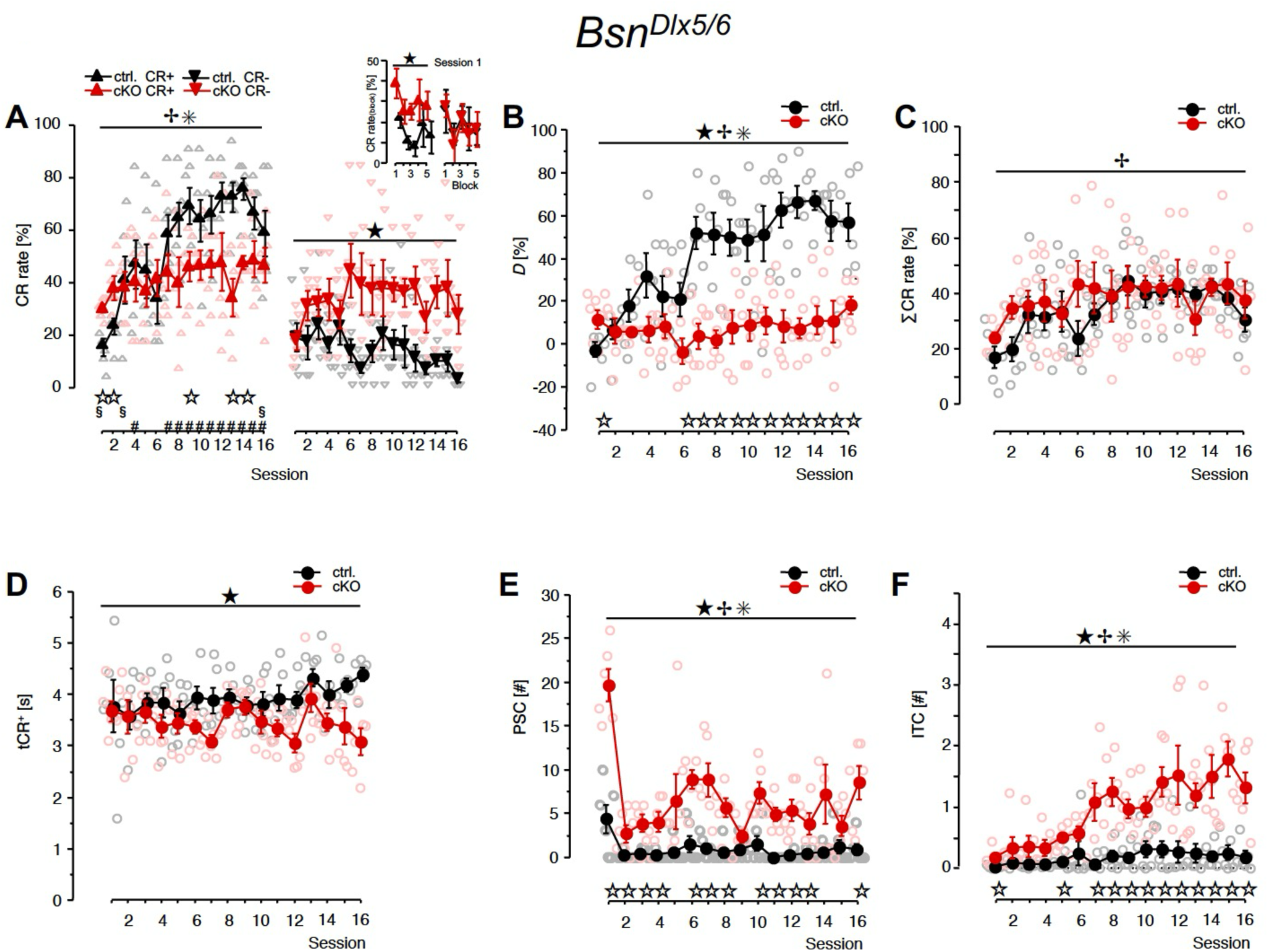
Behavioral characteristics of mice with conditional *Bsn* ablations in a set of GABAergic neurons. Data were collected in a two-way shuttle box used for training of *Bsn^Dlx5/6^*cKO mice (cKO, n=6) and corresponding control mice (ctrl., n=6) on the FM discrimination task once a day for 16 sessions in total. **A,** Relative frequencies of correct conditioned responses (CR+) and false alarms (CR-) per training session. *Inset:* Session 1 was subdivided into five blocks of 12 trials; shown are the relative frequencies of CR+ and CR- per trial block. **B,** Discrimination rate (*D*), *i.e.*, the difference between the relative frequencies of CR+ and CR- per session. **C,** Total frequency of hurdle crossings in response to FMs (ΣCR), *i.e.*, the sum of CR+ and CR− expressed as per cent of the total number of trials per session. **D,** Avoidance latency (tCR+), *i.e.,* the average time to initiate CR+. **E,** Pre-session activity (PSC), *i.e.*, the number of hurdle crossings per 3-min pre-session adaptation period. **F,** Inter-trial activity (ITC), *i.e.*, the average number of hurdle crossings per inter-trial interval. Filled symbols: group means ± S.E.M. Empty symbols: individual data points. ^★^Significant effect of genotype, ^✢^significant effect of session, ^✳^ significant genotype x session interaction (*P*<0.05, RM-ANOVA). ^⋆^Significant difference between corresponding values of ctrl. and cKO; ^#^ctrl.: CR+ rate significantly different from the corresponding CR- rate; ^§^cKO: CR+ rate significantly different from the corresponding CR- rate (*P*<0.05, *t*-test).

In *Bsn^Emx1^* cKO mice (Fig. 1G-L) CBF increased in secondary motor cortex (Fig.1G), primary motor cortex and adjacent anterior cingulate cortex (Fig. 1J, K), somatosensory cortex (Fig. 1K), visual cortex (Fig. 1L) and in parts of the superior colliculus (Fig. 1L). CBF decreased in parts of the olfactory bulb (Fig. 1G), the dorsal peduncular cortex / dorsal tenia tecta region (Fig, 1H), in different patches within the dorsal striatum (Fig. 1I, K) and in the region of the posterior cortical amygdala and adjacent piriform cortex (Fig. 1L).

Compared to the alterations in *Bsn^Emx1^* and *Bsn^Dlx5/6^*cKO mice, the changes in CBF in *Bsn^DAT^* cKO mice were small with respect to both, the magnitude of the changes and the affected areas in the brain (Fig. 1M to R). CBF increased in orbitofrontal cortex (Fig. 1M), in a patch within the dorsal striatum (Fig. 1N), in the dorsomedial hypothalamus (Fig. 1O) and in the brainstem raphe region (Fig. 1R). CBF decreased in parts of the hippocampal formation peaking over the dentate gyrus (Fig. 1P, Q) and in the presubiculum (Fig. 1R).

As major differences in basic brain activity patterns between the various *Bsn* cKOs and corresponding controls occurred in cortical areas, we tested the mice in a learning paradigm that involves a functional auditory cortex, the FM discrimination paradigm [49].

### Data analysis across FM discrimination experiments

First, we compared the three genetic mouse variants that are wild-type for *Bsn* and served as controls in the FM discrimination experiments for differences in behavioral phenotypes. These are *Bsn^lx/lx^* mice (reference for *Bsn^Dlx5/6^*inhibitory cKO, n=6; and for *Bsn^Emx1^* excitatory cKO, n=6), *Bsn^+/+^* x *Emx1^tm1(cre)Krj^*mice (additional reference for *Bsn^Emx1^* cKO, n=6), and *Bsn*^+/+^ x *Slc6a3^tm1.1(cre)BKmn^* mice (reference for *Bsn^DAT^*cKO, n=12) (cf. Additional file 1: Table 1). To this end, data collected in the shuttle box experiments were analyzed using a 3 x 16 (variant x training session) RM-ANOVA (Additional file 2: Table S2). Behavioral changes occurring during the course of the learning experiment were manifested in significant main effects of training session on the CR+ rate, CR- rate, and PSC. Importantly, neither main effects of variant nor variant x training session interaction effects became evident, suggesting that the genetic variants of mice that are wild-type for *Bsn* showed similar FM discrimination learning and performance. For reasons of clarity, these mice are hereafter uniformly referred to as controls. For an initial assessment of behavioral effects of *Bsn* gene ablation at different types of forebrain synapses, data of the control mice and cKO mice acquired in the shuttle box were compared across all the three series of FM discrimination experiments of the present study using a 3 x 2 x 16 (experimental series x genotype x training session) RM-ANOVA. The resulting statistical data (Additional file 3: Table S3A) indicate significant main effects and/or interaction effects of the factors ‘experimental series’, ‘genotype’, and ‘training session’ on the values of dependent variables. To elucidate the sources of the interactions, data were first analyzed separately within the populations of mice with control genotype and cKO genotype using a 3 x 16 (experimental series x training session) RM-ANOVA. Significant main effects of training session on the values of dependent variables were revealed in each genotype (Additional file 3: Tables S3B and S3C), indicating that the controls as well as the cKO mice showed behavioral changes over sessions. Within the control genotype (Additional file 3: Table S3B), main effects of experimental series became evident for only one of the dependent variables (*i.e.*, tCR+), which might point out behavioral differences amongst the independent sets of animals used in the sequentially performed series of FM discrimination experiments and/or some variation in experimental parameters due to, *e.g*., seasonal influences. Importantly, experimental series x session interaction effects did not reach the level of statistical significance. This suggests that the sets of mice used as controls in the three sequentially performed series of experiments showed similar temporal patterns of behavioral changes during learning. In contrast, within the cKO genotype, highly significant main effects of experimental series as well as experimental series x session interaction effects were disclosed (Additional file 3: Table S3C). With the understanding that – as measured by the learning performance of the controls – the three series of FM discrimination experiments were run under comparable conditions, these findings point out major differences in learning and behavior in the shuttle box amongst the three groups of cKO mice under study.

To assess potential relationships amongst the variables monitored in the shuttle box, linear correlation analysis was performed (Additional file 4: Table S4). The rates of CR+ and CR- were not linearly related to each other and were differentially related to PSC and ITC, implying that the behavioral responses to the conditioned stimuli did not simply reflect levels of arousal and activity.

To summarize, the overall pattern of results collected in the FM discrimination experiments is consistent with the hypothesis that changes in the brain caused by conditional absence of Bassoon from sets of GABAergic, glutamatergic, or dopaminergic synapses had distinctive impacts on FM discrimination learning and performance in the shuttle box. For a more detailed examination, data were analyzed separately within each series of FM discrimination experiments.

### Effects of conditional ablation of *Bsn* in sets of GABAergic forebrain interneurons on FM discrimination learning

We next evaluated FM discrimination learning in *Bsn^Dlx5/6^*cKO mice, which lack Bassoon expression primarily, but not exclusively, in PV+ interneurons. Figure 2A displays the mean rates of CR+ (left panel) and CR- (right panel) monitored per training session in *Bsn^Dlx5/6^* cKO and respective control mice. Analysis of the CR+ rate using a 2 x 16 (genotype x training session) RM-ANOVA revealed no significant genotype effect (*F*_1,10_=3.804, *P*=0.0797), a significant session effect (*F*_15,150_=8.756, *P*<0.0001) and a significant genotype x session interaction (*F*_15,150_=3.907, *P*<0.0001). When RM-ANOVA was performed within the control genotype, a significant session effect became evident (*F*_15,75_=11.641, *P*<0.0001), indicating an increasing CR+ rate over sessions. The *Bsn^Dlx5/6^* cKO mice started the experiment with significantly more hurdle crossings than the controls in response to the rising FM used as CS+ (session 1: *t*_10_=3.938, *P*=0.0028; session 2: *t*_10_=2.535, *P*=0.0296; *t*-test). During the course of the experiment, however, an increase in the CR+ rate of *Bsn^Dlx5/6^* cKO mice, if any, was weak and did not reach statistical significance (session effect: *F*_15,75_=1.067, *P*=0.4009; RM-ANOVA). Consequently, the CR+ rate of the control mice rose above the values of the mutants later in the experiment (session 9: *t*_10_=-2.558, *P*=0.0285; session 13: *t*_10_=-4.152, *P*=0.0020; session 14: *t*_10_=-5.812, *P*=0.0002; *t*-test). RM-ANOVA comparing the CR- rate over sessions across genotypes revealed a significant genotype effect (*F*_1,10_=17.980, *P*=0.0017), indicating that *Bsn^Dlx5/6^* cKO mice performed in general more false alarms than the control group. Changes in the CR- rates over sessions as well as genotype differences in the temporal patterns of such changes did not reach statistical significance (session effect: *F*_15,150_=1.003, *P*=0.4549; genotype x session: *F*_15,150_=1.345, *P*=0.1826).

To examine whether the somewhat unexpected preferred responding to that FM that was used as CS+ was innate to *Bsn^Dlx5/6^* cKO mice or acquired during the course of session 1, data collected in session 1 were subdivided into five blocks of trials. The mean rates of CR+ and CR- calculated per trial block of session 1 for controls and *Bsn^Dlx5/6^* cKO mice are shown in the inset of Fig. 2A. Analysis of the CR+ rate using a 2 x 5 (genotype x trial block) RM-ANOVA, with trial block serving as the repeated measure, revealed as expected a significant genotype effect (*F*_1,10_=15.505, *P*=0.0028). However, no effect of trial block (*F*_4,40_=1.339, *P*=0.2722) and, importantly, no genotype x trial block interaction became evident (*F*_4,40_=0.057, *P*=0.9938). These findings suggest that *Bsn^Dlx5/6^* cKO mice performed more hurdle crossings in response to CS+ than the control group throughout session 1, starting with trial block 1. In contrast, the CR-rates recorded in *Bsn^Dlx5/6^* cKO mice and controls during the course of session 1 were very similar (genotype effect, *F*_1,10_=0.007, *P*=0.9344; trial block effect, *F*_4,40_=1.799, *P*=0.1481; genotype x trial block, *F*_4,40_=0.157, *P*=0.9588; RM-ANOVA), implying that the genotype effect on the CR+ rate may not simply reflect general effects of *Bsn* ablation on the sensitivity and responsiveness to sounds *per se* or on locomotor activity.

For a more detailed examination of the discrimination performance, the CR+ rates calculated per genotype and training session were compared with the corresponding values of the CR- rate. Within the reference genotype, the CR+ rate did not differ significantly from the CR- rate before session 4 of differential conditioning (Fig. 2A; sessions 1-3: −0.913≤*t*_5_≤2.231, 0.4032≥*P*≥0.0760; session 4: *t*_5_=3.048, *P*=0.0285; *t*-test). This indicates that the control mice started the experiment without preferential behavioral responding to one of the conditioned stimuli. During the course of discrimination training, the CR+ rate of the controls increased over the CR- rate and, starting with session 7, the control mice consistently performed significantly more correct conditioned responses than false alarms per session (sessions 7-16: 3.948≤*t*_5_≤14.670, 0.0109≥*P*≥2.66 x 10^-5^; *t*-test). In contrast, *Bsn^Dlx5/6^* cKO mice performed significantly more CR+ than CR- in isolated sessions scattered throughout the experiment, starting with session 1 (session 1: *t*_5_=2.621, *P*=0.0470; session 3: *t*_5_=5.000, *P*=0.0041; session 16: *t*_5_=4.466, *P*=0.0066; *t*-test).

To quantify the discrimination performance, the mean discrimination rate *D*, that is, the difference between the rates of CR+ and CR- per training session, was calculated for control and cKO mice (Fig. 2B). RM-ANOVA comparing *D* over sessions across genotypes revealed significant effects of genotype (*F*_1,10_=19.960, *P*=0.0012) and session (*F*_15,150_=9.302, *P*<0.0001) and a significant genotype x session interaction (*F*_15,150_=7.520, *P*<0.0001). During session 1, the *D* value of *Bsn^Dlx5/6^* cKO mice significantly exceeded that of the control group (*t_10_*=2.605, *P*=0.0263, *t*-test), reflecting mainly the higher CR+ rate of *Bsn^Dlx5/6^* cKO mice in this session. During the course of the experiment, however, the controls, but not the cKO mice, significantly improved their discrimination scores (session effect: ctrl., *F*_15,75_=16.343, *P*<0.0001; cKO, *F*_15,75_=0.735, *P*=0.7422; RM-ANOVA). Consequently, starting from session 6, the *D* values of the *Bsn^Dlx5/6^* cKO mice fell significantly behind those of the controls (−2.378≤*t*_10_≤-6.771, 0.0388≥*P*≥4.92 x 10^-5^, *t*-test), indicating a deficit of the cKO mice in improving their discrimination behavior.

To assess potential effects of *Bsn* ablation in GABAergic forebrain synapses on learning a sound-evoked hurdle reaction in the shuttle box, the relative frequencies of total hurdle crossings in response to the FMs *per se* (ΣCR, calculated from the sum of the CR+ and CR-per session) were examined. RM-ANOVA comparing ΣCR over sessions across genotypes disclosed a significant session effect (*F*_15,150_=3.216, *P*=0.0001), while neither a genotype effect (*F*_1,10_=0.792, *P*=0.3943) nor a genotype x session interaction (*F*_15,150_=0.989, *P*=0.4692) became evident. Thus, ΣCR of the cKO mice were very similar to the control values and increased over sessions (Fig. 2C), suggesting that the cKO mice learned and performed the hurdle reaction in response to the sounds *per se*.

To look for effects of *Bsn* ablation in sets of GABAergic interneurons on arousal and activity and on sensory, motivational and/or motor mechanisms, the avoidance latencies (tCR+) and the numbers of PSC and ITC were monitored (Fig. 2D-F). RM-ANOVA comparing tCR+ over sessions across genotypes revealed a significant genotype effect (*F*_1,10_=11.261, *P*=0.0073), indicating that the *Bsn^Dlx5/6^* cKO mice showed shorter reaction times than the controls in response to CS+ (Fig. 2D). The session effect (*F*_15,150_=1.034, *P*=0.4241), and the genotype x session interaction (*F*_15,150_=1.311, *P*=0.2022) did not reach statistical significance. Furthermore, RM-ANOVA comparing PSC as well as ITC over sessions across genotypes revealed significant effects of genotype (PSC, *F*_1,10_=43.168, *P*<0.0001; ITC, *F*_1,10_=59.402, *P*<0.0001) and session (PSC, *F*_15,150_=9.610, *P*<0.0001; ITC, *F*_15,150_=5.541, *P*<0.0001) and genotype x session interactions (PSC, *F*_15,150_=3.945, *P*<0,0001; ITC, *F*_15,150_=3.301, *P*<0.0001). *Bsn^Dlx5/6^* cKO mice performed on average more PSC and ITC than the controls. The difference in PSC (Fig. 2E) between cKO and control mice was highest on the day of initial training, when the shuttle box was novel. Subsequently, PSC decreased steeply in both genotypes (session effect: ctrl., *F*_15,75_=2.787, *P*=0.0018; cKO, *F*_15,75_=7.450, *P*<0.0001 RM-ANOVA). In sessions 2 to 16, genotype differences in PSC were less pronounced and occasionally failed to reach the level of statistical significance (session 1: *t*_10_=6.319, *P*<0.0001; sessions 2-4, 6-8, 10-13, 16: 2.493≤*t*_10_≤6.100, 0.0318≥*P*≥0.0001; *t*-test). The ITC of *Bsn^Dlx5/6^* cKO mice (Fig. 2F), though already exceeding the value of controls, was lowest within the first training session. During the course of the experiment, ITC significantly increased in *Bsn^Dlx5/6^* cKO mice but not in the controls (session effect: cKO, *F*_15,75_=4.882, *P*<0.0001; ctrl., *F*_15,75_=1.123, *P*=0.3515; RM-ANOVA). Consequently, starting with session 7, the *Bsn^Dlx5/6^* cKO mice consistently performed significantly more ITC than the control mice (sessions 1, 5, 7-16: 2.276≤*t_10_*≤5.031, 0.0461≥*P*≥0.0005; *t-*test).

To summarize, while starting the differential conditioning experiment with preferential behavioral responses to the rising FM, *Bsn^Dlx5/6^* cKO mice showed a deficit in the improvement of the FM discrimination reaction over sessions. The deficit was characterized by a flatter, non-significant increase in the CR+ rate and a higher CR- rate compared to the control mice. Increasing numbers of CS-evoked responses over sessions suggest that mechanisms enabling an association between the detection of the sounds *per se* and the likelihood of foot shocks are intact. Compared to the controls, *Bsn^Dlx5/6^* cKO mice showed slightly shorter avoidance latencies and, in general, higher numbers of pre-session and inter-trial crossings.

### Effects of conditional *Bsn* ablation in glutamatergic forebrain neurons on FM discrimination learning

The effect of Bassoon deficiency at glutamatergic synapses of excitatory forebrain neurons on FM discrimination learning was tested in *Bsn^Emx1^* cKO mice. Fig. 3A shows the mean CR+ and CR- rates monitored per training session in *Bsn^Emx1^* cKO and respective control mice. Analysis of the CR+ rate with a 2 x 16 (genotype x training session) RM-ANOVA revealed no significant genotype effect (*F*_1,17_=3.775, *P*=0.0688), a significant session effect (*F*_15,255_=23.008, *P*<0.0001) and a significant genotype x session interaction (*F*_15,255_=2.954, *P*=0.0002). A significant session effect on the CR+ rate became evident in each genotype (ctrl., *F*_15,165_=23.377, *P*<0.0001; cKO, *F*_15,90_=6.980, *P*<0.0001; RM-ANOVA), indicating that both the controls and the cKO mice improved their CR+ rate over sessions. Comparisons within sessions disclosed that *Bsn^Emx1^*cKO mice performed significantly more CR+ than the controls in sessions 2 and 4-6 (2.308≤*t*_17_≤3.799, 0.0338≥*P*≥0.0014, *t*-test). RM-ANOVA comparing the CR- rate over sessions across genotypes revealed a significant session effect (*F*_15,255_=6.187, *P*<0.0001) but no genotype effect (*F*_1,17_=0.020, *P*=0.8895), and no genotype x session interaction (*F*_15,255_=0.988, *P*=0.4685), indicating that the CR- rates of cKO mice and controls were very similar and decreased significantly over sessions. Both genotypes started the discrimination training in session 1 without a preference for one of the FMs used as conditioned stimuli. Starting with session 3 and session 2, respectively, controls and *Bsn^Emx1^* cKO mice performed significantly more CR+ than CR- per session (ctrl.: session 1, *t*_11_=1.372, *P*=0.1974; session 3-16, 3.038≤*t*_11_≤13.780, 0.0113≥*P*≥2.77 x 10^-8^; cKO: session 1, *t*_6_=0.000; session 2-16: 3.500≤*t*_6_≤14.717, 0.0128≥*P*≥6.19 x 10^-6^; *t*-test).

**Figure 3.**
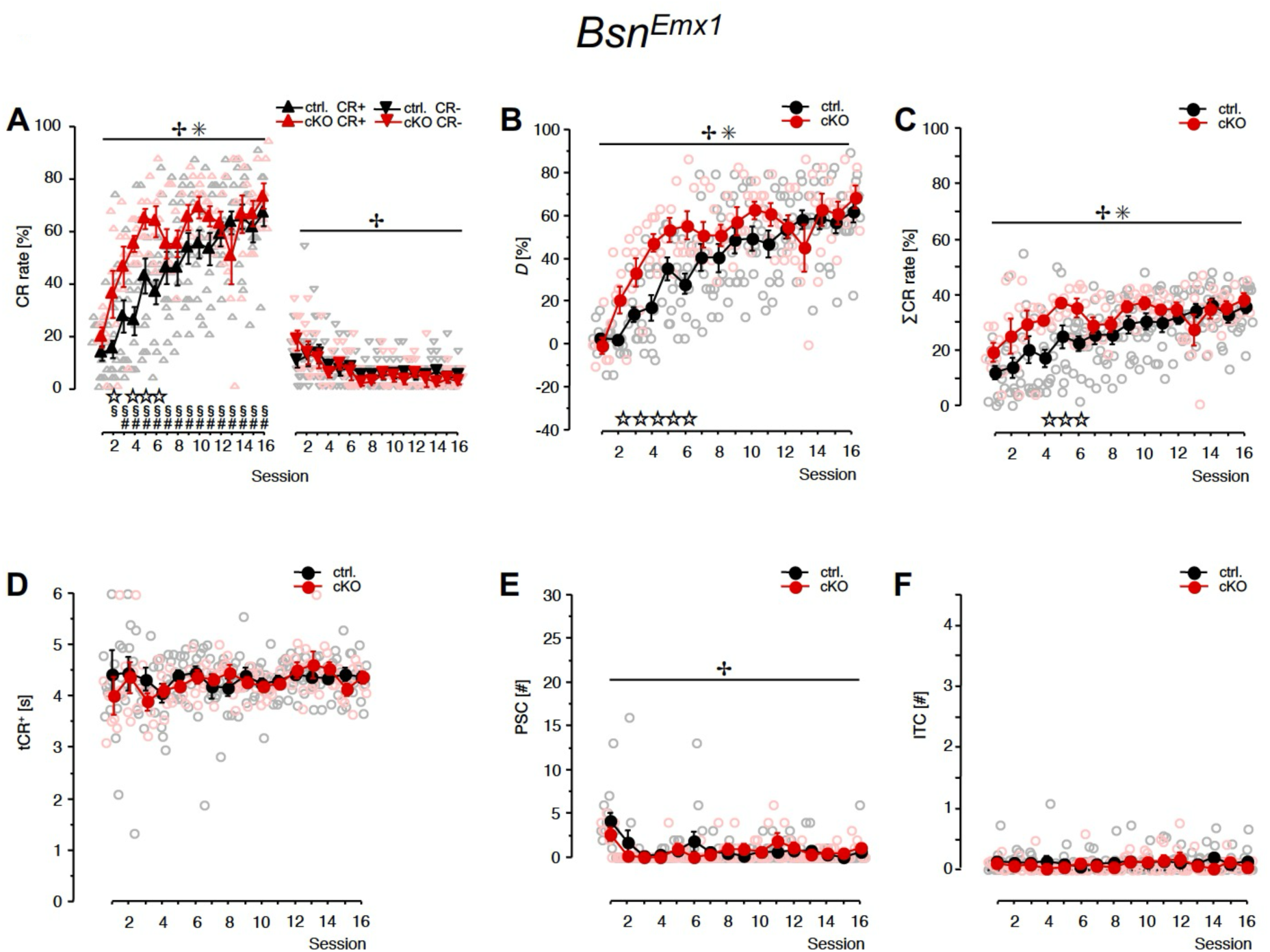
Behavioral characteristics of mice with conditional *Bsn* ablations in glutamatergic neurons. Data were collected in a two-way shuttle box used for training of *Bsn^Emx1^* cKO mice (cKO, n=7) and corresponding control mice (ctrl., n=12) on the FM discrimination task once a day for 16 sessions in total. **A,** Relative frequencies of correct conditioned responses (CR+) and false alarms (CR-) per training session. **B,** Discrimination rate (*D*), *i.e.*, the difference between the relative frequencies of CR+ and CR-per session. **C,** Total frequency of hurdle crossings in response to FMs (ΣCR), *i.e.*, the sum of CR+ and CR− expressed as per cent of the total number of trials per session. **D,** Avoidance latency (tCR+), *i.e.,* the average time to initiate CR+. **E,** Pre-session activity (PSC), *i.e.*, the number of hurdle crossings per 3-min pre-session adaptation period. **F,** Inter-trial activity (ITC), *i.e.*, the average number of hurdle crossings per inter-trial interval. Filled symbols: group means ± S.E.M. Empty symbols: individual data points. ^✢^Significant effect of session, ^✳^significant genotype x session interaction (*P*<0.05, RM-ANOVA). ^⋆^Significant difference between corresponding values of ctrl. and cKO; ^#^ctrl.: CR+ rate significantly different from the corresponding CR- rate; ^§^cKO: CR+ rate significantly different from the corresponding CR- rate (*P*<0.05, *t*-test).

As expected from the curves of the CR+ and CR- rates, the discrimination rates *D* calculated for *Bsn^Emx1^*cKO mice increased faster over sessions than those of controls (Fig. 3B). RM-ANOVA comparing *D* over sessions across genotypes disclosed no significant genotype effect (*F*_1,17_=3.501, *P*=0.0786), a significant session effect (*F*_15,255_=39.032, *P*<0.0001) and a significant genotype x session interaction (*F*_15,255_=3.642, *P*<0.0001). Highly significant main effects of session became evident on the *D* values of both cKO mice (*F*_15,90_=13.671, *P*<0.0001) and controls (*F*_15,165_=34.031, *P*<0.0001), indicating that the mice of each genotype improved their discrimination scores over sessions. Comparisons within each session disclosed that *Bsn^Emx1^* cKO mice reached significantly higher discrimination scores than controls in sessions 2-6 (2.238≤*t*_17_≤3.726, 0.0389≥*P*≥0.0017, *t*-test).

Fig. 3C shows the rates of total hurdle crossings in response to the FM tones *per se* for cKO mice and controls. RM-ANOVA comparing ΣCR over sessions across genotypes revealed no significant genotype effect (*F*_1,17_=3.423, *P*=0.0817), a significant session effect (*F*_15,255_=9.278, *P*<0.0001), and a significant genotype x session interaction (*F*_15,255_=2.017, *P*=0.0147). While both genotypes showed increasing rates of hurdle crossings over sessions (session effect: ctrl., *F*_15,165_=10.891, *P*<0.0001; cKO, *F*_15,90_=2.664, *P*=0.0022), *Bsn^Emx1^* cKO mice performed significantly more hurdle crossings than controls in sessions 4-6 (2.184≤*t*_17_≤3.340, 0.0433≥*P*≥0.0039, *t*-test), reflecting the higher CR+ rates monitored in the group of cKO mice during these sessions.

Avoidance latencies, PSC and ITC are shown in Fig. 3D-F for *Bsn^Emx1^* cKO mice and controls. RM-ANOVA disclosed no significant differences between genotypes (tCR+: genotype effect, *F*_1,17_=0.251, *P*=0.6228; session effect, *F*_15,255_=0.735, *P*=0.7478; genotype x session, *F*_15,255_=0.558, *P*=0.9051. PSC: genotype effect, *F*_1,17_=0.063, *P*=0.8055; session effect, *F*_15,255_=4.184, *P*<0.0001; genotype x session, *F*_15,255_=1.306, *P*=0.1985. ITC: genotype effect, *F*_1,17_=0.305, *P*=0.5877; session effect, *F*_15,255_=0.636, *P*=0.8440; genotype x session, *F*_15,255_=0.916, *P*=0.5473).

To summarize, mice lacking Bassoon at glutamatergic forebrain synapses seem to learn the task faster than controls at the ascending part of the learning curve, but reach similar success rates after the 6^th^ session.

### Effects of conditional *Bsn* ablation in dopaminergic neurons on FM discrimination learning

Finally, we tested mice lacking Bassoon at dopaminergic release sites (*Bsn^DAT^* cKO). As shown in Fig. 4A, the mean relative frequencies of CR+ (left panel) as well as CR- (right panel) monitored per training session were very similar for *Bsn^DAT^* cKO and control mice. Accordingly, RM-ANOVA over sessions across genotypes revealed significant session effects (CR+: *F*_15,330_=32.134, *P*<0.0001; CR-: *F*_15,330_=2.493, *P*=0.0017), but no genotype effects (CR+: *F*_1,22_=0.633, *P*=0.4347; CR-: *F*_1,22_=0.054, *P*=0.8184) and no genotype x session interaction effects (CR+: *F*_15,330_=0.614, *P*=0.8635; CR-: *F*_15,330_=0.499, *P*=0.9405). Both genotypes started discrimination training in session 1 without a preference for one of the FMs used as conditioned stimuli but subsequently performed significantly more CR+ than CR- (ctrl.: session 1, *t*_11_=-0.287, *P*=0.7795; session 2-16, 3.909≤*t*_11_≤17.195, 0.0024≥*P*≥2.69 x 10^-9^; cKO: session 1, *t*_11_=- 1.694; *P*=0.1184; session 2-16: 4.294≤*t*_11_≤15.543, 0.0013≥*P*≥7.83 x 10^-9^; *t*-test).

**Figure 4.**
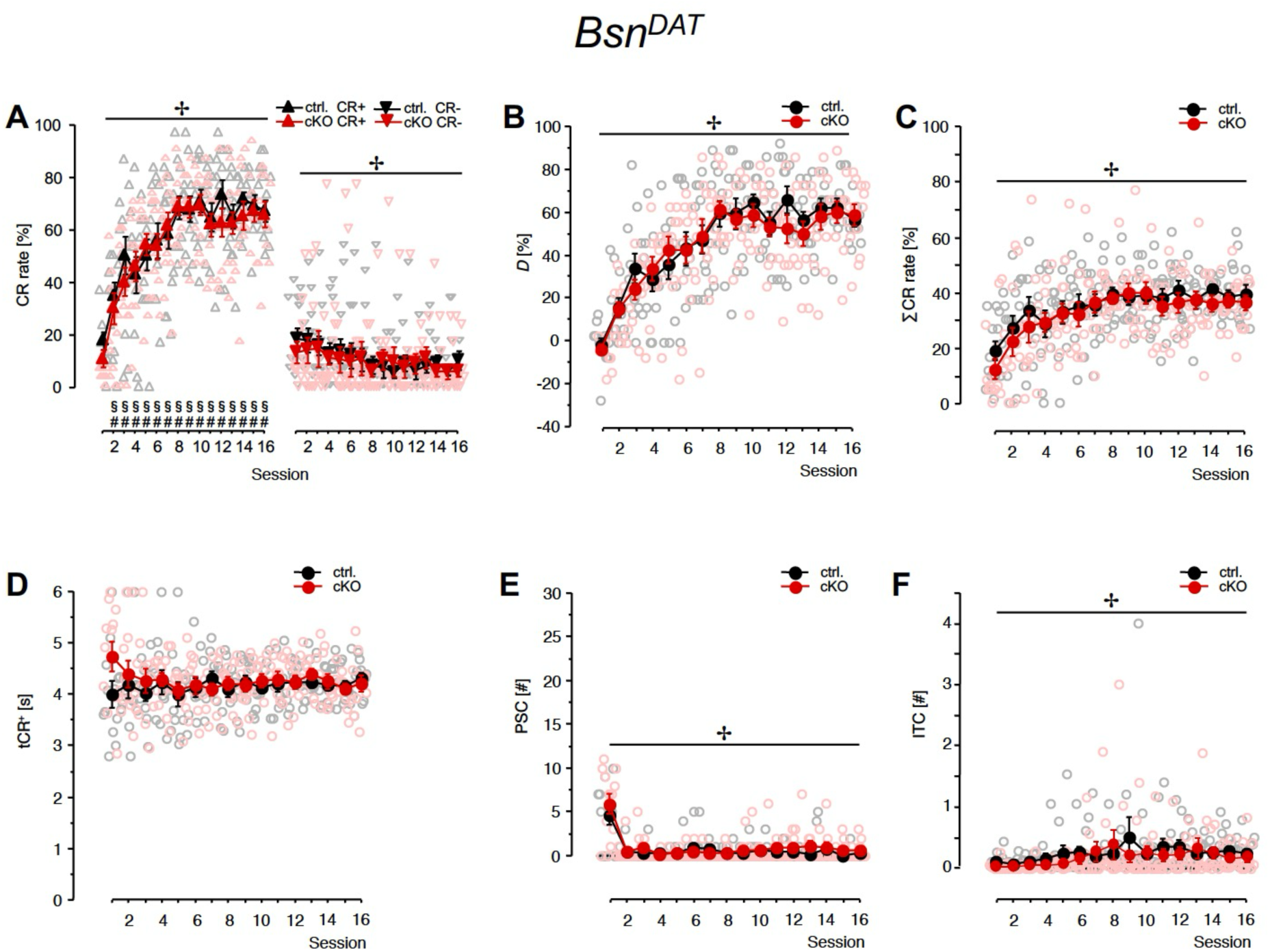
Behavioral characteristics of mice with conditional *Bsn* ablations in dopaminergic neurons. Data were collected in a two-way shuttle box used for training of *Bsn^DAT^* cKO mice (cKO, n=12) and corresponding control mice (ctrl., n=12) on the FM discrimination task once a day for 16 sessions in total. **A,** Relative frequencies of correct conditioned responses (CR+) and false alarms (CR-) per training session. **B,** Discrimination rate (*D*), *i.e.*, the difference between the relative frequencies of CR+ and CR-per session. **C,** Total frequency of hurdle crossings in response to FMs (ΣCR), *i.e.*, the sum of CR+ and CR− expressed as per cent of the total number of trials per session. **D,** Avoidance latency (tCR+), *i.e.,* the average time to initiate CR+. **E,** Pre-session activity (PSC), *i.e.*, the number of hurdle crossings per 3-min pre-session adaptation period. **F,** Inter-trial activity (ITC), *i.e.*, the average number of hurdle crossings per inter-trial interval. Filled symbols: group means ± S.E.M. Empty symbols: individual data points. ^✢^Significant effect of session (*P*<0.05, RM-ANOVA). ^#^Ctrl.: CR+ rate significantly different from the corresponding CR- rate; ^§^cKO: CR+ rate significantly different from the corresponding CR-rate (*P*<0.05, *t*-test).

Expectedly, the mean values of *D* (Fig. 4B) and ΣCR (Fig. 4C) were very similar and showed similar increases over sessions for *Bsn^DAT^* cKO mice and controls (*D*: genotype effect, *F*_1,22_=0.265, *P*=0.6117; session effect, *F*_15,330_=38.614, *P*<0.0001; genotype x session, *F*_15,330_=0.713, *P*=0.7714; ΣCR: genotype effect, *F*_1,22_=0.410, *P*=0.5288; session effect, *F*_15,330_=11.511, *P*<0.0001; genotype x session, *F*_15,330_=0.488, *P*=0.9463; RM-ANOVA).

Also, neither tCR+, nor PSC, nor ITC did show any significant genotype differences (Fig. 4D-F: tCR+: genotype effect, *F*_1,22_=0.409, *P*=0.5291; session effect, *F*_15,330_=0.688, *P*=0.7965; genotype x session, *F*_15,330_=1.112, *P*=0.3439. PSC: genotype effect, *F*_1,22_=0.899, *P*=0.3533; session effect, *F*_15,330_=19.980, *P*<0.0001; genotype x session, *F*_15,330_=0.834, *P*=0.6390. ITC: genotype effect, *F*_1,22_=0.294, *P*=0.5930; session effect, *F*_15,330_=2.125, *P*=0.0087; genotype x session, *F*_15,330_=0.722, *P*=0.7621; RM-ANOVA).

In essence, mice lacking Bassoon in DAT-expressing neurons showed a very similar FM discrimination learning performance in the shuttle box as the respective control mice.

## Discussion

Deletion of the active zone component Bassoon affects the performance of different types of brain synapses differentially. Utilizing CBF-SPECT for functional neuroimaging of baseline activation patterns in the awake state we observed strong deactivations in cortical, notably sensory cortical regions, and weak to moderate activations in parts of the dorsal striatum of *Bsn^Dlx5/6^* cKO mice in comparison to corresponding control mice. This correlates with strongly impaired FM discrimination learning: though *Bsn^Dlx5/6^* cKO mice started the differential conditioning experiment with dominant responding to the rising FM used as CS+, deficits in discrimination improvement became manifest during the course of the experiment. Learning and performance of the hurdle reaction in response to the sweeps *per se* was normal. *Bsn^Dlx5/6^*cKO mice showed shorter avoidance latencies and higher pre-session and inter-trial activities than the controls. In contrast to the widespread decrease of cortical CBF in *Bsn^Dlx5/6^* cKO mice, changes in cortical CBF in *Bsn^Emx1^* cKO mice were more complex with a marked decrease in the dorsal peduncular cortex / dorsal tenia tecta region and increases in secondary and primary motor and in sensory cortices. Notable decreases in CBF were found in several regions of the dorsal striatum. *Bsn^Emx1^* cKO mice learned the FM discrimination faster than the controls. In *Bsn^DAT^* cKO mice, mean CBF changes were comparably small, showing some increases in orbitofrontal cortex, striatum, dorsomedial hypothalamus and in the brainstem raphe. No obvious deficits in discrimination learning were observed in these mice.

### Differential effects of synapse-specific *Bsn* ablation on brain activity

The alterations in cortical CBF in *Bsn^Dlx5/6^* and *Bsn^Emx1^* cKO mice were not accompanied by significant changes in CBF in thalamic nuclei. It seems likely, therefore, that the cortical activations or deactivations are not due to disturbed thalamic inputs, but to either shifts in the intracortical excitation/inhibition (E/I) balance and/or alterations in cortico-striatal circuits. At first glance, the fact that weakened intracortical GABAergic synaptic transmission in *Bsn^Dlx5/6^* cKO mice should result in decreased cortical CBF, while weakened intracortical glutamatergic drive in *Bsn^Emx1^* cKO mice would result in increased cortical CBF, at least in some regions, may seem paradoxical. Both effects, however, could be explained by alterations in cortical inhibition or disinhibition. Cortical PV+ and vasoactive intestinal peptide-expressing interneurons, for instance, are well-known for their crucial role in disinhibitory control [62, 63]. Amongst others, these neurons mediate dopamine actions [64], are activated by reinforcement signals [62, 65] and contribute to associative learning [66]. Network disturbances with disproportionately strong decreases in synaptic transmission of these neurons could thus contribute to both, the decrease in CBF and the poor performance in FM discrimination experiments monitored in *Bsn^Dlx5/6^*cKO mice. Conversely, an increased disinhibition may enhance the CBF in *Bsn^Emx1^*cKO mice and potentially promote performance.

Likely, the complex alterations of the baseline activation patterns in *Bsn^Dlx5/6^* and *Bsn^Emx1^* cKO mice cannot be attributed to dominant effects on disinhibition alone. The reduced CBF in the dorsal peduncular cortex / dorsal tenia tecta region and the posterior cortical amygdala in *Bsn^Emx1^* cKO mice, for instance, do not fit into this scheme. The detailed mechanisms behind these network activation changes remain to be determined.

## What might underly the strong effect of Bassoon deficiency at GABAergic synapses?

In stark contrast to *Bsn^Emx1^* cKO and *Bsn^DAT^* cKO mice, *Bsn^Dlx5/6^* cKO mice showed substantial deficits in FM discrimination learning. Previous investigations on primary hippocampal neuronal cultures and on hippocampal slices revealed commonalities and differences of Bassoon ablation on synaptic functions. In mice lacking functional Bassoon at all synapses excitatory synapses seem to have reduced sizes of synaptic vesicle pools and decreased synaptic strength (*e.g.*, [18, 67]) whereas no significant change is observed at inhibitory synapses in primary hippocampal cultures [67]. Yet, global *Bsn* mutants are suffering from strong rapidly generalizing epileptic seizures [18, 19, 21] suggesting a massive disturbance of the E/I balance in the mutant brains. In hippocampal primary neurons derived from mice lacking Bassoon only at GABAergic synapses (*Bsn^Dlx5/6^* cKO) synaptic vesicle cycling at inhibitory synapses as well as the frequency of miniature inhibitory postsynaptic potentials in ventral hippocampal slices are significantly reduced [19, 46]. Nonetheless, a remarkable reduction of basic transmission at Schaffer collateral-CA1 synapses in the ventral hippocampus is observed and again points to complex network effects and disturbed E/I balance of *Bsn* ablation in GABAergic interneurons [46]. Several behavioral changes monitored in the aforementioned and in the present study resemble those reported by other labs after experimental interference with inhibitory systems in different species, including effects on locomotor activity, novelty seeking, fear, reaction time for conditioned motor responses, and auditory discrimination behavior [68–74].

### Effects of conditional *Bsn* ablation in distinct GABAergic interneurons

FM discrimination learning in the shuttle box involves an instrumental conditioning, in which the association of two conditioned stimuli (tone sweeps) with the meanings ‘Go’ and ‘No-Go’, respectively, are deduced from the successful avoidance of a foot shock (for review see [49]). Initially, there is an association between the detection of the sounds and the likelihood of foot shocks as reinforcers. With the increasing chance to avoid foot shocks by crossing the hurdle during CS+ presentation and by suppressing this response during CS- presentation, these meaningful associations are formed and must be recalled in order to select the appropriate response strategy.

In the present study, the rate of hurdle crossings in response to the FMs *per se* and its increment during the course of the learning experiment was normal in *Bsn^Dlx5/6^* cKO mice when compared to the controls. This indicates that the detrimental effect of the genetic manipulation on discrimination learning was not caused by actions on general mechanisms that may interfere with auditory reinforcement learning and performance in the shuttle box, such as the sensitivity of sensory and motor systems and associative mechanisms required for CS detection and for acquisition and performance of the hurdle reaction to avoid foot-shock. In line with these observations, we did not find significant alterations in CBF in subcortical nuclei providing auditory input to the auditory cortex.

Rising and falling FMs elicit discernible spatial patterns of activity in the auditory cortex, topographically organized in part on the basis of spectro-temporal tuning properties of single neurons in cortical maps [75–77]. Contrast-shaping inhibitory mechanisms in the auditory cortex enhance auditory discrimination [78, 79](for review, see [49, 80]). The severe deficit in FM discrimination learning observed in *Bsn^Dlx5/6^* cKO mice could be caused by detrimental effects of *Bsn* ablation in GABAergic synapses on cortical tuning properties and, as a consequence, impaired auditory discrimination abilities. However, in mutant mice with chronically reduced cortical inhibition [81], narrower spectral tuning rather implies contrast sharpening within the auditory cortex. Assuming comparable alterations after *Bsn* ablation in GABAergic interneurons, naïve *Bsn^Dlx5/6^* cKO mice would be expected to show normal or even better auditory discrimination abilities than controls. Indeed, *Bsn^Dlx5/6^* cKO mice, but not the controls, performed significantly more hurdle crossings in response to the rising FM as compared to the falling one at the beginning of differential conditioning. This indicates that *Bsn^Dlx5/6^* cKO mice are able to perform the sensory processing required to discriminate between the conditioned stimuli as well as the sensorimotor integration and behavior necessary to demonstrate the discrimination reaction. These findings support the view that inhibition in the auditory cortex may control not only learned but also unconditioned innate behaviors that rely on auditory discrimination [68].

Auditory discrimination learning is thought to induce plastic rearrangements of auditory-cortical connections facilitating the sensitive discrimination of the auditory stimuli as well as the integration of auditory stimulus processing with non-auditory cognitive functions, thus enabling meaningful associations and triggering appropriate motor responses. In the present study, the control mice acquired the discriminative behavior during differential conditioning to the FMs and subsequently improved it mainly by increasing the rate of responses to the rising FM used as the Go-stimulus CS+. In contrast, the *Bsn^Dlx5/6^* cKO mice – although preferentially responding to the rising FM at the beginning of differential conditioning – failed to increase the CR+ rate over sessions. This deficit in improvement of the Go-response may point, at least in part, to effects of the *Bsn* ablation on the integration of already processed auditory information with multisensory and non-sensory information required for the behavioral manifestation of the discriminative memory [49, 82], such as the association of the conditioned stimuli with the respective Go and No-Go meanings, decision making and appropriate response selection in order to achieve the goal of the discrimination task.

Goal-directed learning involves NMDA-type glutamate receptor-mediated cortico-striatal connections [83]. In auditory Go/No-Go tasks, the auditory cortex may contribute to sensorimotor associations and response selection during goal-directed behavior [84, 85]. Accordingly, after sound discrimination learning the expression of synaptic NMDA receptors is differentially regulated in auditory-cortical and striatal regions [86, 87]. Moreover, the functional interaction of the auditory cortex with striatal regions increases selectively when the auditory stimulus is associated with goal-directed behavior, that is, during encoding the transformation of the Go-stimulus representation into the appropriate motor response [88], and long-term memory required to express this goal-directed behavior crucially depends on proper NMDA receptor-mediated excitatory signaling within the auditory cortex [51]. In mutant mice with chronically reduced cortical inhibition, a down-regulation of auditory-cortical excitatory drive was reported [81]. Assuming similar effects of *Bsn* ablations in inhibitory neurons, the abolished increment in the Go-response rate monitored in *Bsn^Dlx5/6^* cKO mice can be explained by postulating severe disturbances of excitatory signaling-dependent mechanisms of FM discrimination learning that support the strengthening of functional connections required for goal-directed behavior.

In goal-directed tasks, successful action selection also critically involves mechanisms to inhibit inappropriate motor responses, such as responses to a No-Go stimulus and premature responses during inter-trial intervals. Impaired cortical GABA transmission has been implicated in high levels of impulsivity, that is, tending to react prematurely, with short response latencies [89, 90]. In auditory discrimination tasks, GABA signaling in auditory cortices – probably in communication with fronto-striatal circuits – plays a role in the inhibitory control of inadequate motor responses in order to attain the current goals [84, 85, 91–93].

*Bsn^Dlx5/6^* cKO mice showed actually shorter response latencies and higher CR- rates, pre-session activities and inter-trial activities than the controls. Given that conditional ablation of *Bsn* in inhibitory forebrain neurons attenuates cortical GABAergic transmission [19], impaired inhibitory control would explain the higher rates of CR- and ITC monitored in *Bsn^Dlx5/6^* cKO mice compared to the controls. Consistent with this view, open field test behavior [45, 46] and the exceptionally high pre-session activity (present study) monitored on the initial day of training, when the maze was novel, indicate a novelty-induced hyperlocomotion phenotype of *Bsn^Dlx5/6^* cKO mice. High novelty reactivity was shown to be predictive for a lack of behavioral inhibition, a type of behavioral impulsivity [94, 95]. Moreover, *Bsn^Dlx5/6^* cKO mice demonstrate an anxiety phenotype [45, 46]. Aside from a genetic background associated with high impulsiveness, ITC in a shuttle box may relate to the fear state of the animals in the context of the maze [96]. Animals increase inter-trial responding after trials in which a shock was applied [97] and may even show “pseudo-learning” by adopting a high level of locomotor activity to minimize exposure to the shock [98]. Consistent with these suggestions, *Bsn^Dlx5/6^*cKO mice increased the ITC significantly during the course of the experiment. The positive correlation between the rates of CR- and ITC monitored in the present study (*cf.* Additional file 4: Table S4) may point to common factors – such as the impulsiveness and/or the fear state of the animals – in the control of these behavioral parameters.

Goal-directed action control involves a broad cortical-basal ganglia network including, besides others, sensory, retrosplenial, parietal, orbital, cingulate and motor cortices, striatum and amygdala [99]. The present neuroimaging study performed on *Bsn^Dlx5/6^* cKO mice revealed significant decreases in mean CBF in auditory, visual, retrosplenial, cingulate and motor cortices and in the amygdala as well as increases in striatum when compared to control mice. It seems likely that these widespread network changes are related or contribute to changes in goal-directed action.

### Effects of conditional *Bsn* ablation in glutamatergic forebrain neurons

In *Bsn^Emx1^* cKO mice lacking Bassoon at glutamatergic forebrain synapses, prominent increases in CBF were found relative to controls in some of the cortical regions that had demonstrated decreases in *Bsn^Dlx5/6^* cKO mice, including sensory, retrosplenial, cingulate and motor cortices. In the striatum, CBF was decreased relative to controls, again opposite to the pattern seen in *Bsn^Dlx5/6^*. These mutants exhibited a significantly faster increase in the Go-response rate than the controls at the ascending part of the FM discrimination learning curve. By analogy with the abolished increment in the Go-response rate monitored in *Bsn^Dlx5/6^*cKO mice, the accelerated increment in the Go-response rate monitored in *Bsn^Emx1^*cKO mice can be explained by postulating improved excitatory signaling-dependent processes of FM discrimination learning that support the strengthening of functional connections required for goal-directed behavior. This is in line with a previous study showing enhanced learning in hippocampus-dependent tasks of *Bsn^Emx1^* cKO, due to the juvenile phenotype of the dentate gyrus [44]. While the hippocampus is likely not required for active avoidance and FM discrimination learning in the shuttle box [100–102], glutamatergic synapses of *Bsn^Emx1^*cKO mice that preserve a more juvenile character might influence brain circuitries and functions relevant for FM discrimination learning.

The SPECT data collected in *Bsn^Dlx5/6^* cKO and *Bsn^Emx1^* cKO mice suggest that Bsn ablation in inhibitory and excitatory forebrain synapses might have opposite effects on mechanisms controlling, amongst others, basal synaptic functions and background excitatory neurotransmission in the naïve, resting animal, which eventually govern neuronal excitability and network activity during goal-directed learning.

### Effects of conditional Bsn ablation in dopaminergic neurons

Cortical dopamine is instrumental in modulating the efficiency of FM discrimination learning and long-term memory formation in rodents [50, 52, 60, 103] and the presence of Bassoon at dopaminergic release sites has been reported [4, 5, 7, 104]. In the present study, however, mice lacking *Bsn* in DAT-expressing neurons showed only moderate alterations in their brain activation patterns and learning curves very similar to those of the controls. Reasons for these unanticipated findings could be that DAT promoter-controlled expression of Cre might not take place in dopaminergic cells relevant for the observed learning behavior [105] or that Bassoon ablation affects dopamine signaling modes that are dispensable for learning the FM discrimination [106].

## Conclusions

The absence of the active zone-organizing protein Bassoon from excitatory forebrain synapses vs. its absence from GABAergic synapses of interneurons, which depend on expression of *Dlx5/6* gene regulatory elements, has quite distinct effects on brain activity and FM-discrimination learning involving the auditory cortex. The absence of Bassoon from release sites of DAT-expressing dopaminergic neurons has a surprisingly weak effect on overall brain activity and FM-discrimination learning performance. A simple correlation of synaptic phenotypes, i.e. basic functions and plasticity, with behavioral outcome is not possible, rather disturbance of network functions and impairment of E/I balance need to be taken in account.

## Abbreviations

CBF: cerebral blood flow
cKO: conditional knockout
CR+: correct conditioned response
CR-: incorrect response (=false alarm)
CS: conditioned stimulus; (CS+, 4-8 kHz; CS-, 8-4 kHz)
E/I: excitation/inhibition
FM: frequency-modulated tone
ITC: inter-trial hurdle crossings
PSC: pre-session hurdle crossings
PV+: parvalbumin-expressing
RM-ANOVA: repeated-measures analysis of variance
SPECT: single-photon emission computed tomography
^99m^Tc-HMPAO: ^99m^Technetium hexamethyl propylene amine oxime
tCR+: avoidance latency

## Acknowledgements

We are very grateful to Chris Theuerkauf, Isabel Herbert, Kristin Marquardt and Holger Reim for expert technical assistance and to the colleagues from LIN animal facility for steady support.

## Author contributions

Conceptualization and design of the study, writing the original version: E.D.G., J.G., W.T. SPECT-CBF experiments and analysis: A.M.O., J.G.

Behavioral experiments: H.S.

Analysis of behavioral experiments: W.T.

Mouse genetics, breeding and basic characterization of mice: A.A., A.F., C.M.-V.

All authors contributed to discussion and editing the final version.

## Supplementary Information

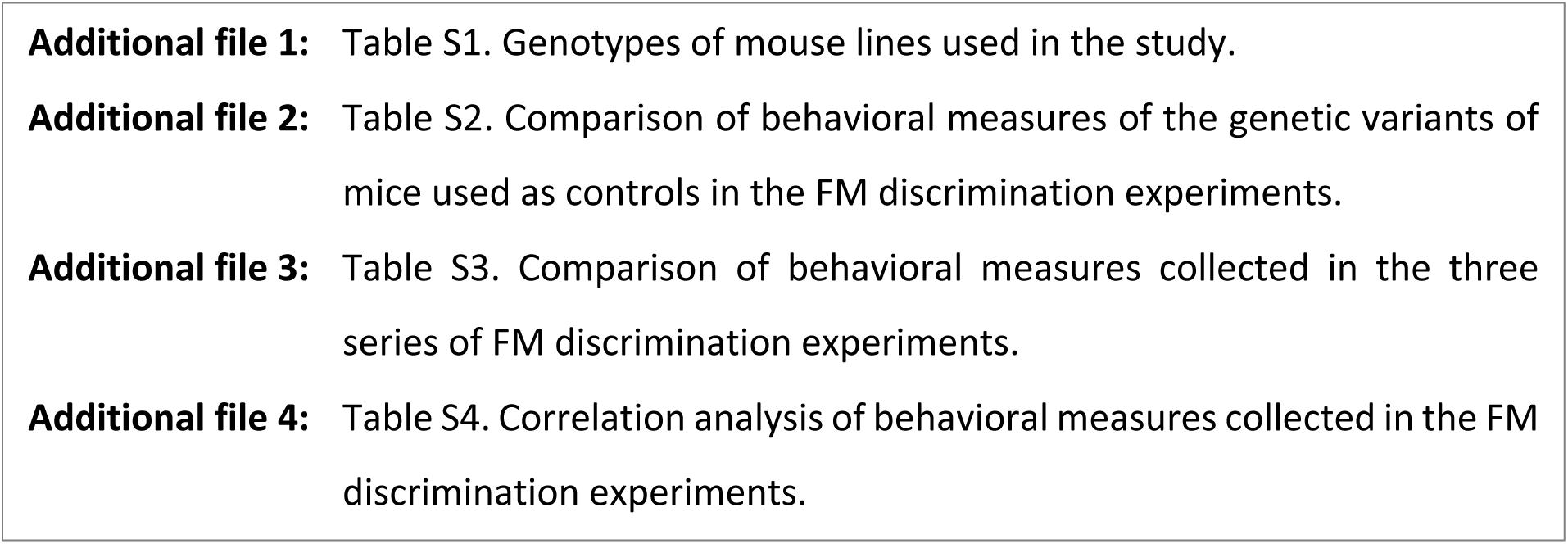

