## Supplementary Tables for "Role of Bassoon-mediated active zone integrity at different types of brain synapses for brain activity and cortex-dependent memory formation"

**Additional Table S1** Genotypes of mouse lines used in the study

| **Short name of cKO** | **Genetics** | **Corresponding controls** |
|---|---|---|
| ***Bsn$^{Dlx5/6}$*** | *Bsn$^{lx/lx}$* X *Tg$^{(dlx5a\text{-}cre)1Mekk/J}$* | *Bsn$^{lx/lx}$ (=Bsn$^{tm1.1Arte}$)* |
| ***Bsn$^{Emx1}$*** | *Bsn$^{lx/lx}$* X *Emx1$^{tm1(cre)Krj}$* | *Bsn$^{lx/lx}$ (=Bsn$^{tm1.1Arte}$)*<br>*Bsn$^{+/+}$* X *Emx1$^{tm1(cre)Krj}$* |
| ***Bsn$^{DAT}$*** | *Bsn$^{lx/lx}$* X *Slc6a3$^{tm1.1(cre)BKmn}$* | *Bsn$^{+/+}$* X *Slc6a3$^{tm1.1(cre)BKmn}$* |

*Bsn$^{lx/lx}$*, mouse line with second exon of the *Bassoon* gene flanked by loxP sites [19]; all other strains are described in detail on The Jackson Laboratory websites: https://www.jax.org/jax-mice-and-services/find-mice

**Additional Table S2.** Comparison of behavioral measures of the genetic variants of mice used as controls in the FM discrimination experiments.

| | Main effect of variant | | Main effect of session | | Variant x session | |
|---|---|---|---|---|---|---|
| | $F_{2,27}$ | $P$ | $F_{15,405}$ | $P$ | $F_{30,405}$ | $P$ |
| CR+ | 2.251 | .1247 | 39.630 | **<.0001** | 1.270 | .1588 |
| CR– | 0.378 | .6886 | 4.957 | **<.0001** | 0.571 | .9683 |
| PSC | 0.815 | .4532 | 11.029 | **<.0001** | 0.886 | .6427 |
| ITC | 1.889 | .1706 | 1.057 | .3959 | 0.648 | .9255 |
| tCR+ | 0.606 | .5529 | 0.533 | .9220 | 0.391 | .9986 |

Shown are values of a 3 x 16 (variant x training session) RM-ANOVA, with training session serving as the repeated measure, comparing behavioral data of *Bsn$^{lx/lx}$* mice (n=12), *Bsn$^{+/+}$* X *Emx1$^{tm1(cre)Krj}$* mice (n=6), and *Bsn$^{+/+}$* X *Slc6a3$^{tm1.1(cre)BKmn}$* mice (n=12). Significant values ($P$<0.05) in bold. CR+, correct conditioned response rate; CR–, false alarm rate; PSC, pre-session activity; ITC, intertrial activity; tCR+, avoidance latency.

**Additional Table S3.** Comparison of behavioral measures collected in the three series of FM discrimination experiments.

| ***A*** (total) | Main effect of exptl. series | | Main effect of genotype | | Exptl. series x genotype | | Main effect of session | | Exptl. series x session | | Genotype x session | | Exptl. series x genotype x session | |
|---|---|---|---|---|---|---|---|---|---|---|---|---|---|---|
| | $F_{2,49}$ | $P$ | $F_{1,49}$ | $P$ | $F_{2,49}$ | $P$ | $F_{15,735}$ | $P$ | $F_{30,735}$ | $P$ | $F_{15,735}$ | $P$ | $F_{30,735}$ | $P$ |
| CR+ | 2.865 | .0666 | 0.255 | .6161 | 4.221 | **.0204** | 51.833 | **<.0001** | 1.755 | **.0080** | 4.523 | **<.0001** | 1.919 | **.0024** |
| CR– | 17.096 | **<.0001** | 7.951 | **.0069** | 7.594 | **.0013** | 3.526 | **<.0001** | 1.383 | .0849 | 1.195 | .2699 | 1.404 | .0756 |
| PSC | 45.006 | **<.0001** | 51.775 | **<.0001** | 39.758 | **<.0001** | 31.544 | **<.0001** | 4.787 | **<.0001** | 4.238 | **<.0001** | 4.407 | **<.0001** |
| ITC | 24.718 | **<.0001** | 21.085 | **<.0001** | 24.021 | **<.0001** | 8.973 | **<.0001** | 3.630 | **<.0001** | 3.499 | **<.0001** | 2.880 | **<.0001** |
| tCR+ | 18.347 | **<.0001** | 3.056 | .0867 | 3.787 | **.0295** | 1.214 | .2548 | 0.708 | .8774 | 1.001 | .4523 | 1.017 | .4419 |

| ***B*** (ctrl.) | Main effect of exptl. series | | Main effect of session | | Exptl. series x session | |
|---|---|---|---|---|---|---|
| | $F_{2,27}$ | $P$ | $F_{15,405}$ | $P$ | $F_{30,405}$ | $P$ |
| CR+ | 3.329 | .0510 | 42.594 | **<.0001** | 1.458 | .0593 |
| CR– | 2.735 | .0829 | 5.274 | **<.0001** | 0.995 | .4763 |
| PSC | 0.345 | .7052 | 11.267 | **<.0001** | 0.588 | .9608 |
| ITC | 1.743 | .1942 | 1.565 | .0803 | 0.819 | .7412 |
| tCR+ | 3.485 | **.0450** | 0.834 | .6396 | 0.564 | .9709 |

| ***C*** (cKO) | Main effect of exptl. series | | Main effect of session | | Exptl. series x session | |
|---|---|---|---|---|---|---|
| | $F_{2,22}$ | $P$ | $F_{15,330}$ | $P$ | $F_{30,330}$ | $P$ |
| CR+ | 5.155 | **.0146** | 16.196 | **<.0001** | 2.261 | **.0003** |
| CR– | 16.407 | **<.0001** | 0.827 | .6477 | 1.478 | .0547 |
| PSC | 53.498 | **<.0001** | 19.512 | **<.0001** | 6.262 | **<.0001** |
| ITC | 40.734 | **<.0001** | 7.857 | **<.0001** | 4.270 | **<.0001** |
| tCR+ | 22.551 | **<.0001** | 1.492 | .1060 | 1.244 | .1820 |

Shown are values of RM-ANOVA, with training session serving as the repeated measure, comparing behavioral data (*A*) of all experimental mice using a 3 x 2 x 16 (experimental series x genotype x training session) design, and of control mice (*B*) or cKO mice (*C*) using a 3 x 16 (experimental series x training session) design. Significant values ($P$<0.05) in bold. CR+, correct conditioned response rate; CR, false alarm rate; PSC, pre-session activity; ITC, intertrial activity; tCR+, avoidance latency.

**Additional Table S4.** Correlation analysis of behavioral measures collected in the FM discrimination experiments.

| | Correlation | *P* |
|---|---|---|
| CR+, CR- | -,006 | ,8664 |
| CR+, ITC | -,022 | ,5230 |
| CR+, tCR+ | -,164 | <,0001 |
| CR+, PSC | -,235 | <,0001 |
| PSC, ITC | ,258 | <,0001 |
| tCR+, PSC | -,275 | <,0001 |
| tCR+, ITC | -,327 | <,0001 |
| CR-, PSC | ,337 | <,0001 |
| CR-, tCR+ | -,442 | <,0001 |
| CR-, ITC | ,505 | <,0001 |

Correlation coefficients (sorted by absolute value) and probability levels (Fisher's r to z transformation) against the null hypothesis that the correlation is equal to zero. 880 observations were used. CR+, correct conditioned response rate; CR-, false alarm rate; PSC, pre-session activity; ITC, intertrial activity; tCR+, avoidance latency.
